# MCFBM: a behavioral analysis system enabling objective inference of songbirds’ attention during social interactions

**DOI:** 10.1101/2023.12.22.573152

**Authors:** Mizuki Fujibayashi, Kentaro Abe

**Author notes:** Corresponding author Kentaro Abe, Ph. D. Professor, Lab of Brain Development, Graduate School of Life Sciences, Tohoku University. Katahira 2-1-1, Aoba-ku, Sendai, Miyagi 980-8577, Japan.

## Abstract

Understanding animal behavior is crucial in behavioral neuroscience, which aims to unravel the mechanism driving these behaviors. A milestone in this field is the analysis of behavioral reactions among animals engaging in social interactions. Although many studies have revealed the fundamental roles of social interaction in social learning, the behavioral aspects of these interactions remain poorly understood, largely due to the lack of tools for analyzing complex behaviors and the attention of subjects in naturalistic, free-moving conditions. Here, we introduce a high-precision system for behavior analysis in songbirds using a marker-based motion capture technique. This system accurately tracks the body location and head direction of freely moving finches and is applicable to multiple subjects during social interaction. With this system, we have quantitatively analyzed behaviors of zebra finches (*Taeniopygia guttata*) related to visual attention. Our analysis revealed variations in the use of right and left eyes, as well as the duration of sight, among the individuals presented. Further analysis and comparison of their behaviors during both virtual and live presentation identified the similarities and differences in their behavioral reactions. Additionally, we observed changes in their behavioral reactions during a conditioned learning paradigm. This system provides an efficient and easy-to-use tool for advanced behavioral analysis in songbirds, providing an objective method to infer their focus of attention.

## Introduction

Given that the primary output of the brain appears on motor control, the analysis of animal behavior has been a central issue of behavioral neuroscience. With the continual development of revolutionary methods to measure the neuronal activity *in vivo* during free moving condition, it has become increasingly important to assay animal behavior with high precision, particularly in a naturalistic setting with minimal behavioral disturbance ^1^. For this purpose, several cutting-edge technologies of marker-less analysis incorporating deep-learning techniques have been devised and utilized ^1,2^. However, even to those methods, achieving efficient, robust and high precision tracking of the trajectory of animal behavior is still a challenging issue ^3^. Specifically, those machine-learning based methods require large datasets for training, and difficult for analyzing the inter-individual behavioral differences, particularly in species with significant variation in body color or shape. In addition, it is still challenging to precisely analyze the behavior of species that exhibits rapid movement in three dimensions (3D) environment such as songbirds, particularly when multiple subjects are located in a single cage. These features are required to analyze in detail the naturalistic interaction of multiple subject to reveal the mechanism underlies inter-individual communication.

Marker based behavioral analysis have been extensively employed for kinematic tracking, especially in humans, capturing highly precise trajectories of specific body parts. Several studies on humans have utilize this precise movement analysis by maker-based motion tracking to discern subject’s attention or emotion ^4–6^. While this approach has been coopted for animal research, it application has largely been confined to larger species ^7^. Marker based tracking holds an advantage over marker-less motion tracking methods due to its ability to precisely track the orientation between the affixed markers ^8^. This feature is especially valuable in the motion tracking of avian species, as they exhibit relatively confined eye movement ^9,10^. By tracking head orientation, it becomes feasible to infer the subject’s focus of attention ^11^. A previous research has employed marker-based head-tracking in pigeons, demonstrating their selective use of specific visual field in response to visual stimuli ^12^. However, the method, employing eight markers on their head and back attached to their feathers, together with wearing backpacks, weighting around 10 g in total, remains impractical for smaller birds like finches.

In this study, we present a maker-based motion tracking system tailored for songbirds. This system provides high-precision tracking of both body location and head direction. Using this technology, we investigated the behavioral response of zebra finches to conspecific recognition in both live and virtual environments. We also examined behavioral changes during experimental conditioning to previously neutral signals. Our methodology offers an efficient and accurate analysis of songbird behaviors.

## Results

### Motion and head direction capture system for small songbirds

To achieve high-precision capture of behavior in free-moving condition consistently across individuals, we adapted and modified a marker-based motion capture system, originally designed for larger animals ^12,13^, for use with small birds. This modified system is henceforth referred to as MCFBM (Motion Capture for Finch Behavior Measurements). In this study, we utilized MCFBM on the zebra finch (*Taeniopygia guttata*), one of the most widely studied model animals to explore the neural and behavioral mechanisms behind animal communication ^14^. MCFBM employs color detection of two light-weight plastic markers, affixed to the head of the subject along with the rostral-caudal axis of midline (Fig. 1a). For ensuring a stable attachment, these markers were attached to a small plastic screw which was firmly cemented to the cranial bone of the subject. The total weight of two marker and screws were 1.26 g. After sufficient acclimation of the screws and markers, the behavioral analyses of subjects were conducted. Differently colored markers were fixed to each screw base (Fig. 1a). The arena for behavioral analysis comprised a metal mesh cage (36.5 × 25.5 × 26 cm) situated within a soundproof chamber (inside dimension: 68 × 50 × 40 cm) (Fig. 1b). Subjects were movie-recorded at 10 frame per seconds (fps) by a single 180° fisheye camera mounted on the chamber’s ceiling. To enhance marker detection accuracy, we replaced the cage’s roof with a clear monofilament net, and removed the perch from the cage in order to increase the accuracy of motion capture in 2D.

**Fig. 1.**
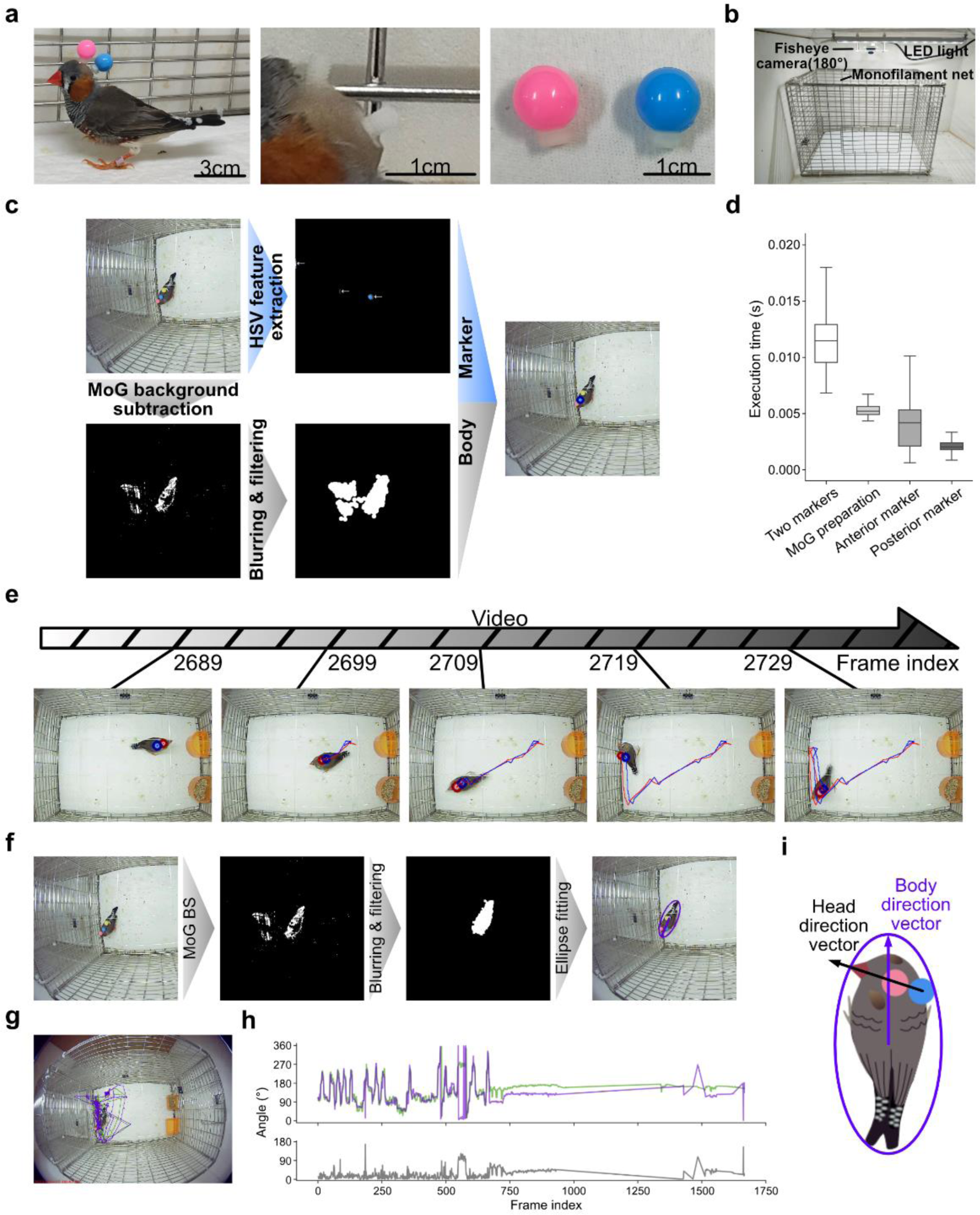
| Configuration of marker-based motion capture system, MCFBM. (**a**) Markers and their attachments. A male zebra finch with markers attached (left), close-up image of screw basements cemented on the cranial bone (middle), and markers (right). (**b**) Arena for monitoring behavior. (**c**) Schematic diagram of the marker tracking procedure. HSV feature extraction (top middle) and MoG background subtraction (MoGBS, bottom left) are applied to the recorded movie (top left), and the results were merged (right). (**d**) Time required for motion analysis with various procedures. The total execution time for two marker detections is composed from MoGBS preparation phase, and subsequent anterior and posterior marker detection phase. Boxplots display the median and the interquartile range. (**e**) Example of marker tracking. Trajectories of two markers over 40 frames (∼ 4 s; frame #2689–2729) are shown in pink and blue for every 10 frames. (**f**) Schematic diagram of body center and body direction estimation. After MoGBS, the body is fitted with ellipse. (**g, h**) Evaluation of background subtraction-mediated approximation of the body center. Trajectory of estimated center (purple) and a marker affixed on the back (green) (**g**), and their traces (**h**). (**h**) Top, a plot showing clockwise angles of body-direction vectors, measured relative to the rightward horizontal vectors. Results from body-direction estimated either by MoGBS-based analysis (purple) and by the attached back marker (green) are shown. Below, angles between body-direction vectors estimated from MoGBS and the back marker. (**i**) Diagram showing the tracked head direction vector (black) and body direction vector (purple).

For color marker tracking, we devised a computer algorithm that utilizes HSV (hue, saturation, value) color feature extraction with the Mixture-of-Gaussians (MoG) background subtraction method (Methods). This combination ensured robust marker detection and body-center detection across various environments, irrespective of the subject’s body color (Figs. 1c,e; Supplementary Movie1). The tracking of markers was shown to be robust. The algorithm detected the anterior head marker in 98.83% of the frames. The measured distance between the two head marker was 9.45 pixels ± 5.82 (mean ± s.d.) throughout the recorded period, the variation of distance reflects the vertical orientation of the head. The algorithm performed robustly across six different experimental setups and among three different birds. The times required for marker detection were 3.86 ms and 6.21 ms for one and two maker extraction, respectively, using HSV-based color feature extraction (Fig. 1d; Methods). The body-center was defined either as the center of an approximated ellipse or as the center of the body area detected by background subtraction (Fig. 1f). We opt for background subtraction methods, because plastic markers cannot be firmly attached to the back of songbirds without obstructing their natural behaviors. Furthermore, we fitted the body area with an ellipse to mitigate fluctuation in body location caused by signal noise. Including the application of MoG background subtraction adds an additional 5.38 ms, thus, a total 11.59 ms is required for the detection of two markers and the body center (Fig. 1d; Methods). To validate the accuracy of body-center tracking using background subtraction methods, we compared it with the accuracy of body-center detection from marker-based tracking, with a plastic marker attached on the subject’s back using adhesives. In this validation, background subtraction-based detection of center of bird were highly correlated with the marker on the back of the subject (Figs. 1g,h). Being confirmed that body center can precisely detected by background subtraction, we finalized our system to be comprised with only two-head makers. Since our marker was stably attached to the bone, we can analyze their behavior repeatedly in a reproductive way.

From the captured location of markers and the features related to the subject’s body, we were able to measure the direction of both their body and head. For this analysis, we utilized one of the four vectors extending from the center of ellipse to its apices, with their predictions based on head movement from prior frames. (Figs. 1f,i). Consequently, in our subsequent analysis using MCFBM, we employed two head markers along with the estimated body center and body direction.

### Female and male directed behavior in zebra finch

To validate our developed MCFBM system, we examined female-directed behaviors in male zebra finch. It is widely recognized that male zebra finches exhibit distinct behavioral response to female presentation: they become notably active, approaching females and engaging in a courtship dance characterized by side-to-side body swaying ^15^. We conducted a marker-based motion capture analysis on male zebra finch during the presentation of male or female conspecifics (Fig. 2a). Upon introducing a female zebra finch adjacent to the male’s cage, the male subject exhibited pronounced activity, often hopping back and forth close to the female cage’s side. This heightened activity was reflected in the significant increase in cumulative moving distance compared to that observed against male birds (Fig. 2b).

**Fig. 2.**
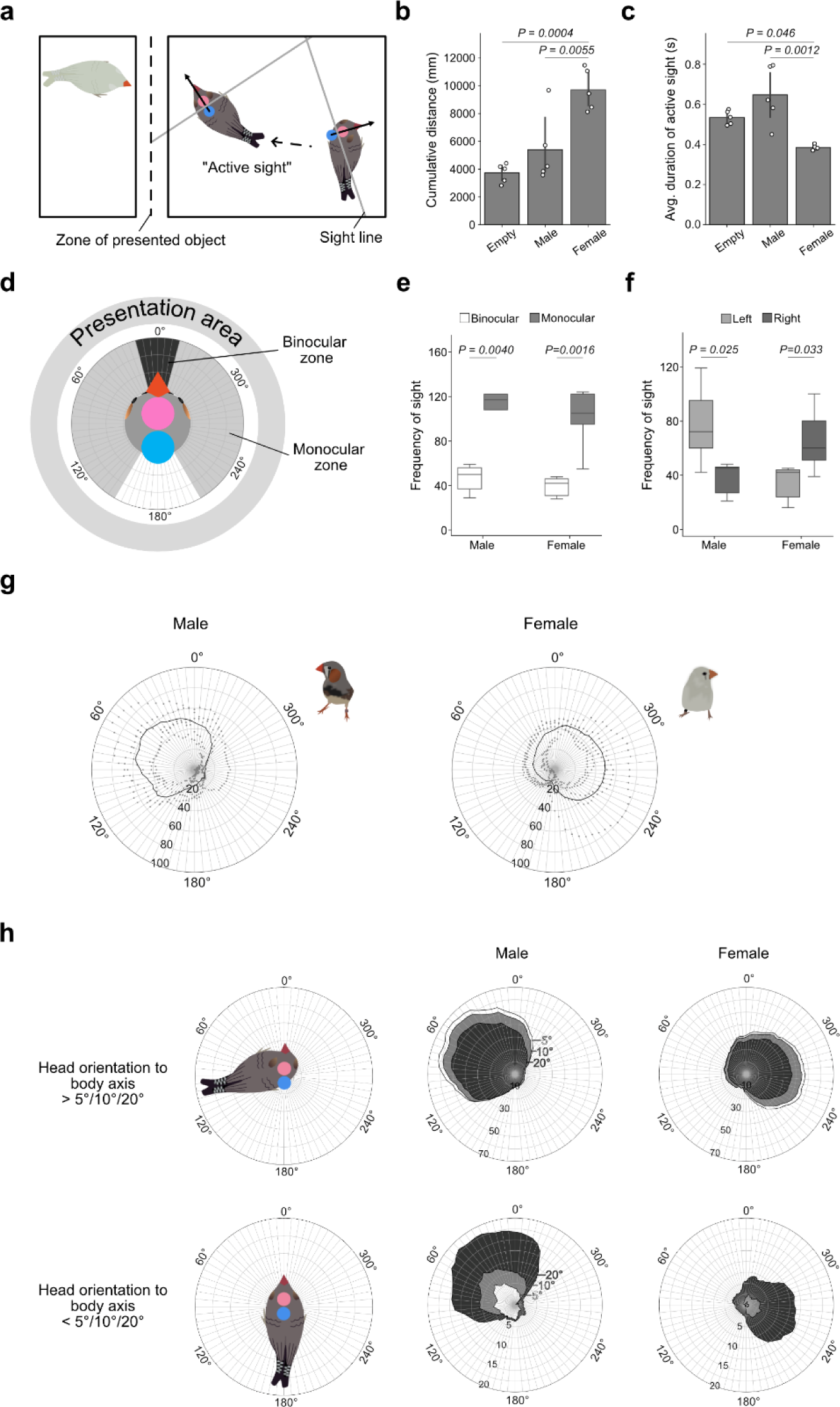
| MCFBM captures social context-dependent behavior in zebra finch and reveals the laterality of eye usage. (**a**) Diagram of experimental setup. We define the ’active sight’ event as the bird movement resulting in the sight line (gray line) crossing the zone of the presented object (broken line), lasting for more than 3 frames. Black arrow, head direction vector. (**b**, **c**) Cumulative moving distances of the bird’s center (**b**) and the average duration of active sight (**c**) in 3-min sessions. Mean ± 95% confident intervals (CI). Tukey HSD test, 5 trials. (**d**) Diagram of visual field usage against presented stimuli. Black and gray fields represent the binocular and monocular fields, respectively. (**e**, **f**) Specific usage of their sight towards male and female targets. Independent-samples *t*-test, 5 trials. Frequency of incidents where the line from presented area persistently projected for more than 3 successive frames within a range beyond 15° in monocular and binocular fields (**e**), and in the left and right eye fields (**f**). Boxplots display the median and the interquartile range ± 95% CI. (**g**) Polar plot showing the frequency of presented area projection lasting for more than 3 frames (approximately 0.3 s), against male (left) and female (right) birds. (**h**) Same as (**g**) differentiating instances where the angle between head-direction and body-direction is either beyond 5°, 10°, 20° (top), or less than 5°, 10°, 20° (bottom).

Visual information is pivotal for social and environmental recognition in songbirds ^16^. Birds are known to orient their heads towards stimuli that capture their attention ^9,17^. Utilizing MCFBM, we aimed to determine if the focus of attention could be inferred from the captured movie. For subsequent sight analysis, we defined the ’sight-line’ as the vertical line bisecting the line connecting the two head markers. This line was used as the basis for analyzing behaviors related to sight (Fig. 2a). To deduce the line of sight, we measured the frequency at which head movements resulted in the sight-line intersecting the zone of the presented object for a duration longer than three frames (approximately 0.3 s). We referred to this occurrence an ’active sight’ event (Fig. 2a). Additionally, we measured the time following the active sight event, which reflect the duration of sight-line remained on the targets without head movements. This measurement was termed ’duration of active sight’. The average ’duration of active sight’ metrics were found to be significantly lower during female presentation compared to male presentation, possibly reflecting the increase in motivated movements (Fig. 2c).

Next, we sought to assess the usage of their sight by analyzing the subject’s head direction. It has been revealed that the visual field of each eye covers about 170° in the horizontal plane in zebra finches ^18^. To deduce their line of sight, we evaluated the angle between two vectors: the frontal head vector, which intersects the two markers on their head, and the vector originating from the midpoint of two markers on their head, directed towards the projection area. The analysis was conducted with a 5° resolution during the less active periods, when the subject was not engaged in extensive back-and-forth hopping. Firstly, we contrasted the usage of binocular and monocular vision by measuring the frequency at which the projection persisted for more than three frames in binocular vison range (± 15° relative to head direction) versus the monocular vision range (± 20°– 150°) (Fig. 2d). We observed a difference in the frequency of monocular versus binocular sight. For observing both males and females, the subject used monocular sight significantly more often than binocular sight (Fig. 2e). Next, we investigated the laterality of their eye usage and found that the usage of the right and left eye depended on the stimuli. The right eye was more often used for observing female birds, whereas the left eye was preferentially used for observing males (Fig. 2f). Further analysis on the visual field usage revealed that the presentation area was often captured in the monocular zone centered around ± 60° relative to the head direction (Fig. 2g). This observation aligns with previous behavioral analysis ^19^, thereby validating that our system can be applied to infer visual attention of songbirds. To further scrutinize behavioral indicators of visual attention, we conducted an additional analysis of eye usage, distinguishing cases based on whether the head was oriented relative to the body axis. This analysis revealed consistent stimulus-dependent laterality in eye usage, which occurred regardless of the head’s orientation along the body axis (Fig. 2h). This suggests that body direction is not a reliable indicator for determining a bird’s visual attention. These observations further indicate that analyzing sight while taking head direction into account, including ’active sight’, is sufficiently reliable for inferring visual attention.

### Marker-based detection of multiple subjects

The detection method of MCFBM utilizes color. Therefore, the number of markers can be increased provided that there is efficient color separation between them. To validate this approach, we conducted marker-based tracking of two subject within a single cage. We introduced either a male or female subject alongside one specific male individual, each affixed with uniquely colored makers. As hypothesized, the same marker detection algorithm used for single subject tracking successfully detected all markers on multiple subjects. However, to accurately determine the center or directional movement of each bird, an additional method to track the silhouettes of each subject was required. Consequently, we integrated Optical Flow ^20^ into our algorithm, in addition to the HSV-based marker detection (Fig. 3a; Supplementary Movie 2). Using these in combination, the marker detection accuracy for individuals during the interaction was 98.64 ± 1.38% (mean ± s.d., *n* = 3 bird). The accuracy of body center prediction was 94.59 % for male and 92.89 % for female, averaging for two subjects, a level sufficiently high to analyze their behavior effectively. Through the analysis using MCFBM, we found that the average distance between two subjects was significantly smaller for male and female pairs than for male and male pairs (Figs. 3b,c). Furthermore, we were able to identify epochs of social interactions between subjects, defined as instances when the two anterior markers of different individuals came within 70 pixels (approximately 87.91 mm) of each other. Notably, such interactions occurred more frequently with females than with males (Figs. 3d,e). These findings suggest that our system is capable of tracking social interactions among multiple subjects within a single enclosure.

**Fig. 3.**
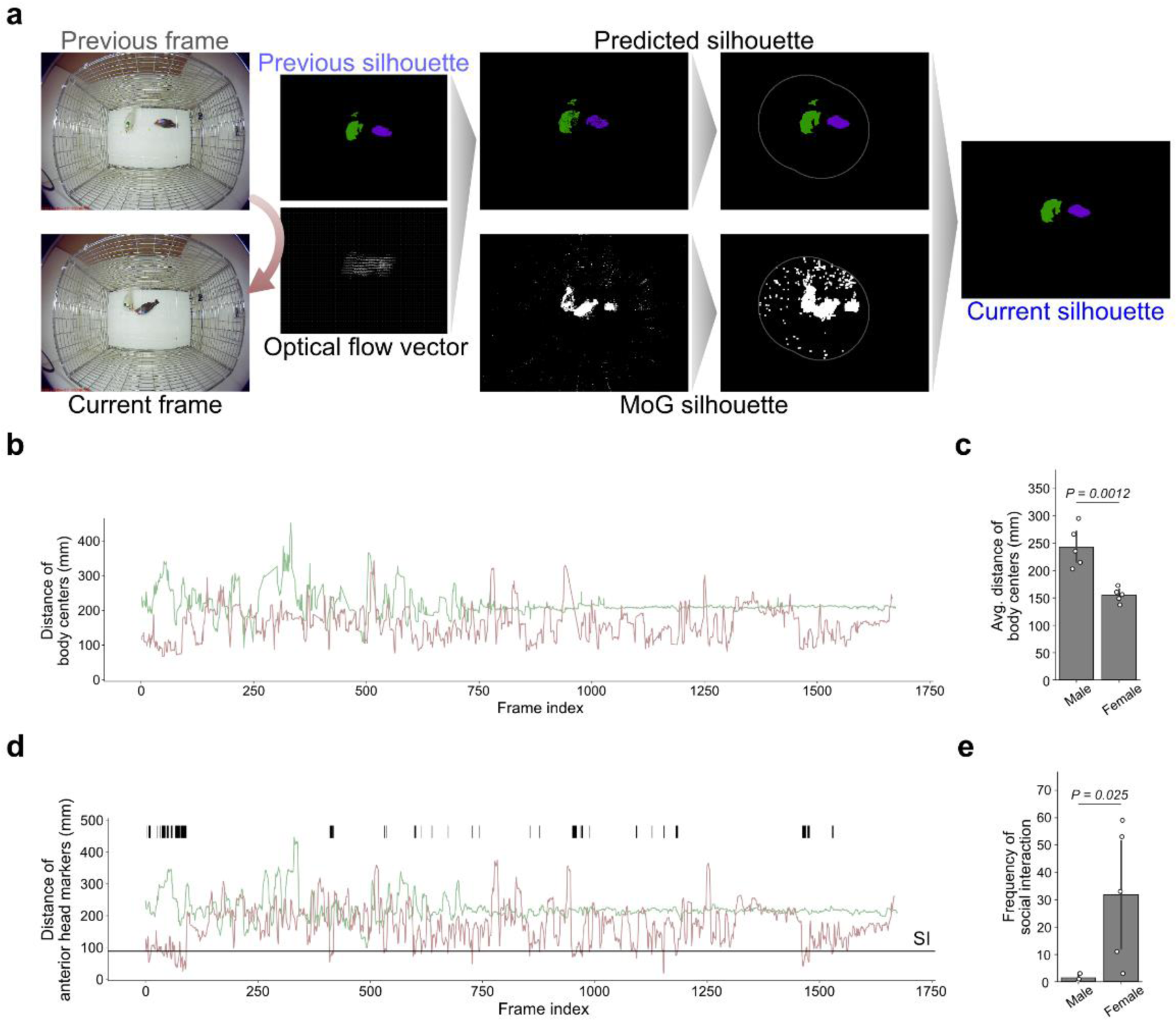
| MCFBM revealed the social interaction of directly interacting multiple subjects. Simultaneous motion capture of two birds interacting in a single cage. (**a**) Schematic diagram for motion capture analysis on multiple subjects. The silhouettes of each bird were identified and used for body location and body direction estimation. (**b**, **c**) Trace plot showing the distance within two birds’ body center during the male-male (green) or male-female (red) interaction for one session (**b**) and the summarized data from five sessions (**c**). Independent-samples *t*-test. (**d**) Trace of the distance between the two anterior head markers on each bird during the male-male (green) or male-female (red) interaction for one session. SI, criterion of social interaction. Raster plot above the trace indicates the detected events of social interaction where the markers were within 70 pixels (∼ 87.90 mm). (**e**) Frequency of social interaction during the 3-min sessions. Bar graphs display the mean ± 95% CI.

### Comparison to live and virtual interaction

Many studies have demonstrated that virtual visual stimuli can effectively substitute for live stimuli in contexts like song learning and inducing courtship behavior ^21,22^. To explore the differences in their response to live versus virtual stimuli, we compared the behavioral reactions to movie-recorded females and males with those elicited by live counterparts. For the virtual stimuli, we presented a movie of birds on an LCD monitor placed adjacent to the subject’s cage and then compared their behavior to that observed during live bird presentation (Fig. 4a). To concentrate on behaviors related to visual stimuli, the movies used in this study did not include audio. The MCFBM revealed both similarities and differences in the behaviors elicited by live and virtual stimuli. Upon the presentation of virtual stimuli, we noted behaviors similar to those observed during live female presentation, such as approaching the monitor and engaging in a characteristic dance-like behavior, involving side-to-side body swaying (Fig. 4b). The ’duration of active sight’ and the ’frequency of gaze’, which was defined as active sight lasting > 1 sec, were reduced for female compared to males in both live and virtual presentation (Figs. 4c,d). The average duration of active sight was comparable between live and virtual stimuli for both male and female (Fig. 4e). However, we also observed a significant decrease in cumulative moving distance in response to the virtual stimuli compared to live presentation in both male and female (Fig. 4f). As observed when live birds are presented, a tendency for increased cumulative distance to virtual female presentation over virtual male presentation was observed, although this difference was not statistically significant (Fig. 4f). These results suggest that male-female discrimination also occurs with virtual information and elicits similar behavioral reactions. However, MCFBM revealed differences in their responses to virtual and live presentations.

**Fig. 4.**
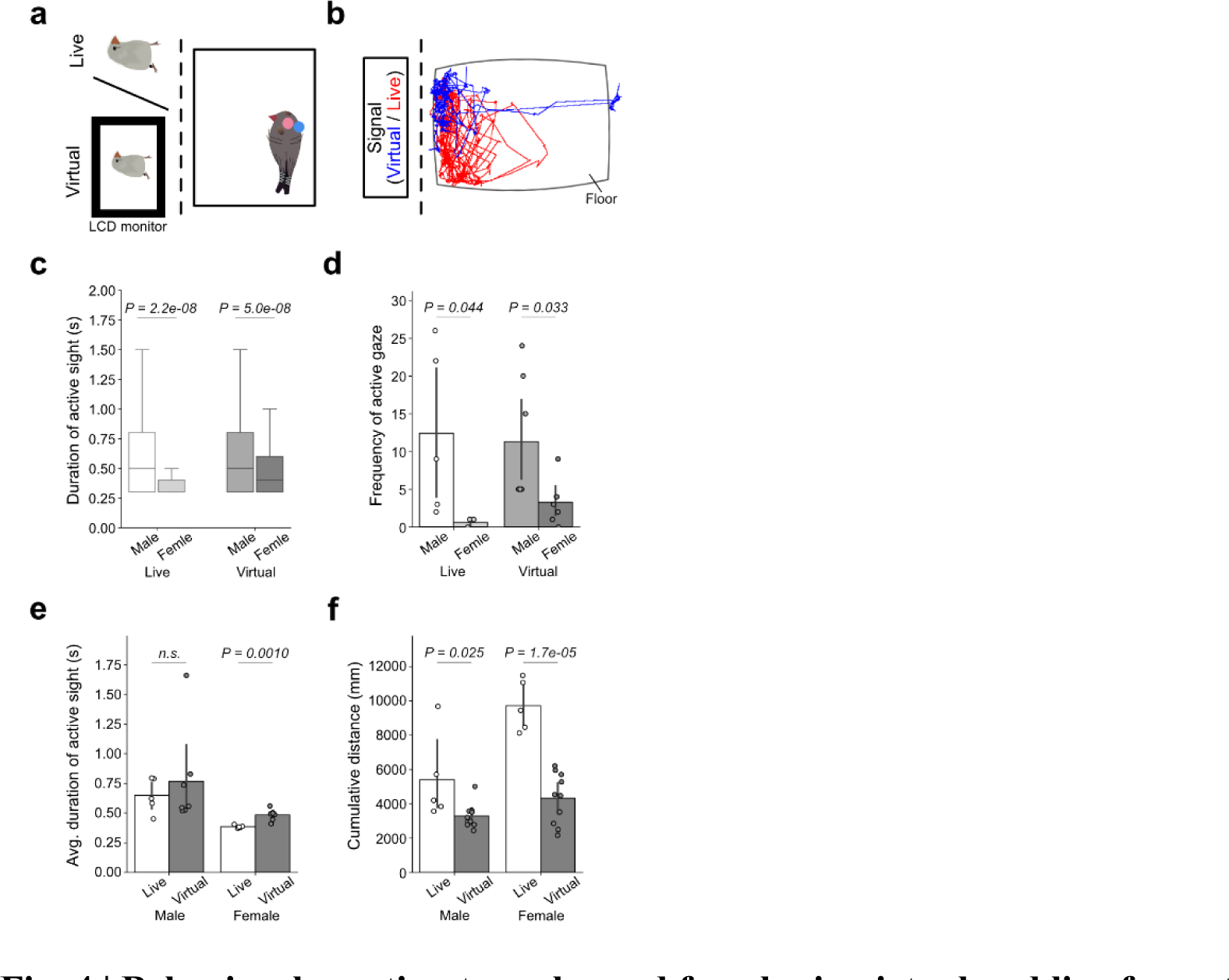
| Behavioral reaction to males and females in virtual and live format. Behavioral response of a male zebra finch to the presentation of other birds in both live and virtual formats. (**a**) Illustration showing the experimental setup of live/virtual bird presentation. (**b**) Traces of the bird’s center during a female presentation in live (red) and virtual format (blue). (**c**) ’Duration of active sight’ across all sessions. Boxplots display the median and the interquartile range ± 95% CI. (**d**) Frequency of ’active gaze’, defined as an active sight event lasting longer than 1 s. (**e**) Average duration of active sight. (**f**) Cumulative distance traveled in 3 min sessions. Independent-samples *t*-test was used for statistical analyses shown in this figure. The number of trials in each cohort: male-live, 5; male-virtual, 7; female-live, 5; female-virtual, 7 in (**c**, **e**); male-live, 355; male-virtual, 409; female-live, 174; female-virtual, 401 in (**c**); male-live, 5; male-virtual, 10; female-live, 5; female-virtual, 10 in (**f**). n.s. not significant. Bar graphs display the mean ± 95% CI.

### Individual discriminations in virtual interactions

Having observed that virtual stimuli can substitute live stimuli in some extent, we proceeded to examine whether virtual stimuli can also convey sufficient information for individual discrimination. We compared behavioral responses to three different cage-mate males and one unfamiliar male, both in live and virtual formats. By analyzing the reaction, we found that certain parameters exhibited correspondence between live and virtual stimuli. Specifically, the average ’duration of active sight’ displayed a positive correlation between live and virtual stimuli among the subjects (Fig. 5a). On the other hand, when deducing their body direction by assessing the percentage of frames in which the presentation area was projected onto the subjects relative to the subject’s body orientation, we observed a reverse trend in directional bias towards the three familiar males (Fig. 5b). Interestingly, this metrics were not correlated to unfamiliar individuals.

**Fig. 5.**
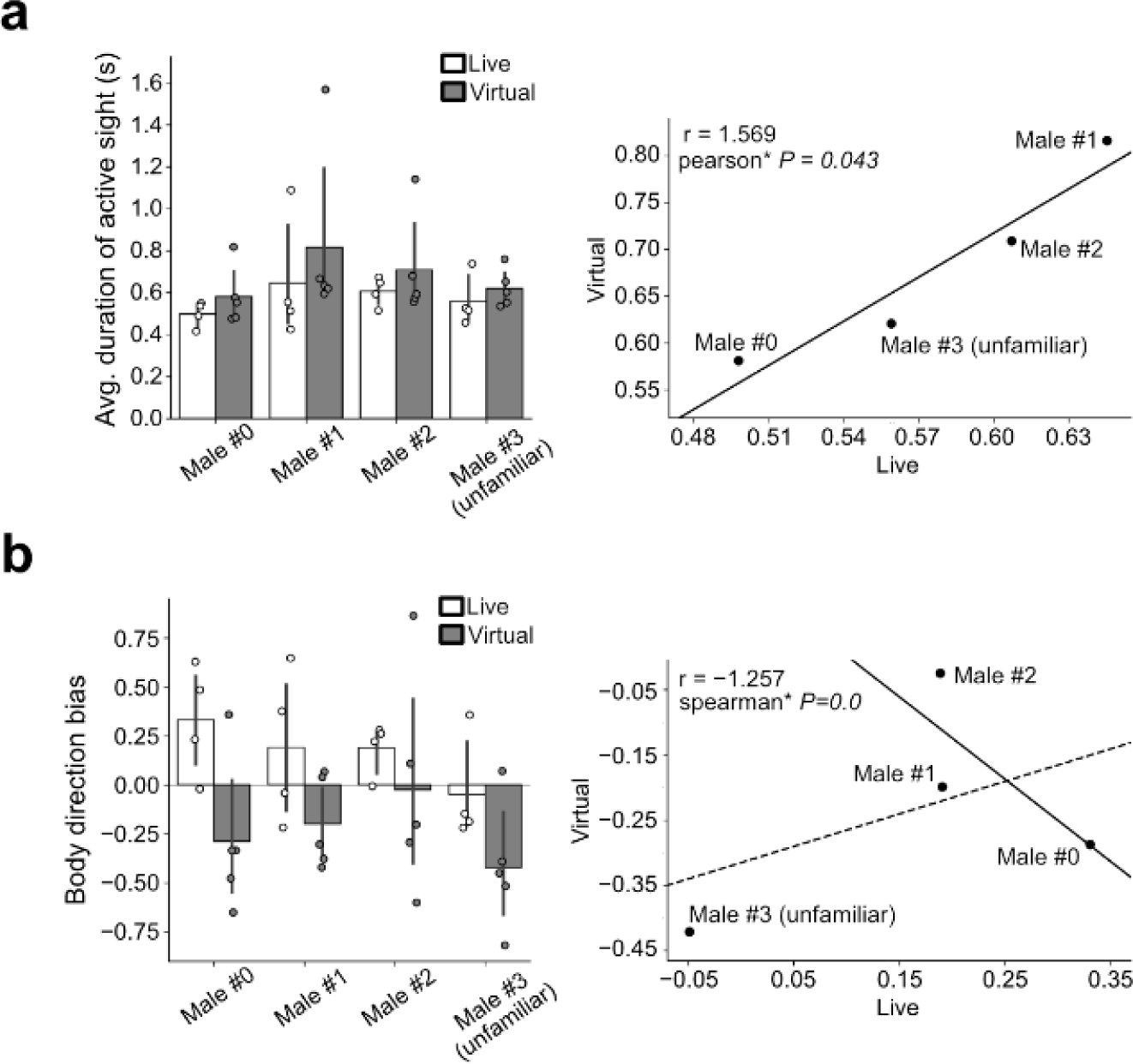
| Virtual stimuli convey sufficient information for individual discrimination. Behavioral responses of a male zebra finch to presentation of identical birds in both virtual and live formats. The results from three familiar males (Males #0, #1, #2) and one unfamiliar male (Male #3) are shown. (**a**) Bar graph showing the duration of active sight (left), and a correlation plot comparing live and virtual presentations (right). The bold line represents the linear regression using the least squares method, with its coefficient of slope and the Pearson correlation *P*-value displayed on the left. (**b**) Same as (**a**), showing the deviation in the ratio of frames where the presented area is projecting forward to the subject while the subject remains still. The bold line represents the linear regression of only the three familiar males, with its coefficient of slope and Spearman’s rank correlation coefficient indicated. The dashed line, the linear regression including all four males. Number of trials in each cohort: live, 4; virtual, 5 trials. Bar graphs display the mean ± 95% CI.

Further, we sought to identify the specific cues facilitating this discrimination. We turned our attention to the role of color information in individual recognition, because color plays an important role in visual communication in zebra finches ^23^. We displayed female movies, wherein the color of subjects was deliberately manipulated (Fig. 6a). We found that certain behavioral parameters were responsive to the effects of color manipulation, while others remained unaffected by it. The average ’duration of active sight’ showed a comparatively difference between the clipped and original movie (Fig. 6b). The difference in ’frequency of sight’, indicated by the instances where the sight-line crossed the monitor, was small between the clipped and original movies, while it was larger between the original and other movies (Fig. 6c). We hypothesized that visual alteration might influence various stages of recognition, leading to complex behavioral responses. Since human visual perception may not always align with how birds recognize objects, analyzing these complex behaviors in response to changes in the visual features of stimuli poses challenges, as a singular parameter may be inadequate to accurately reflect their perception. Hence, a more comprehensive analysis becomes essential for an optimal behavioral assessment. To address this, we employed principle component analysis (PCA) on parameters that responded to changes in live and virtual conspecifics presentations (Fig. 6d). The Euclid distance of initial six principal components from the center of original movie group increased in relation to the degree of color reduction (Fig. 6e). These observations suggest that the behavioral reaction to virtually displayed social information is dependent on the visual features present. Furthermore, our system can be utilize to objectively analyze a bird’s perception of stimuli.

**Fig. 6.**
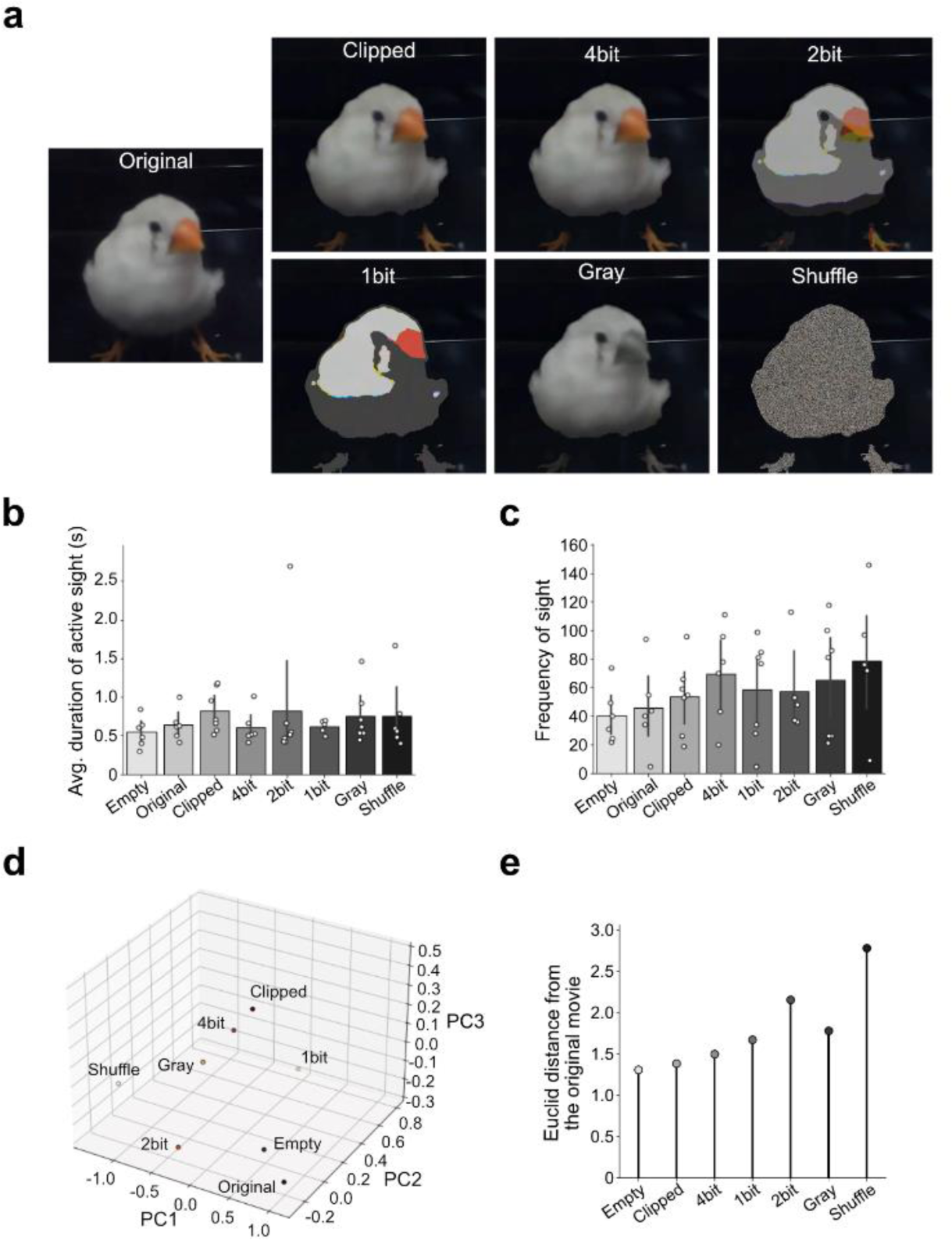
| Color cue of virtual stimuli facilitates individual discrimination. Behavioral response of a single male bird to female movies with altered color features. **(a)** Frames from the presented movies, showcasing the original, clipped, and the color reduced versions. (**b**) Average duration of active sight in response to movie presentation. **(b)** Frequency at which the signal was observed. Number of trials in each cohort: empty, 6; original, 6; clipped, 7; 4-bit, 6; 2bit; 7, 1-bit, 5; gray, 7; shuffle, 6 trials. (**d**, **e**) Principal component analysis of the behavioral reaction. (**d**) Centers for each signal groups represented in the reduced three-dimensional space of the first three principal components (PC1, PC2, and PC3). (**e**) Euclid distances of the initial six principal components from the center of original movie. Each point corresponds to the score of each session in (**b**) and (**c**). Bar graphs display the mean ± 95% CI.

### Assessing the change of behavior during learning

We next turned our attention to understanding how behavior changes in response to the altered salience of the stimuli. We hypothesize that the behavioral reaction to previously neutral information will change following its conditioning with a salient stimulus ^24^. To test this, we presented movie of crows without sound to a naïve bird that was never seen a crow before. For these naïve birds, the crow movie did not elicit a significantly different reaction compared to movies of zebra finch female and Bengalese finch (Figs. 7a,b). To quantify their preference, we analyzed cumulative moving distance and the total number of frames in which the subject was being close to the monitor, defined as instances where the x-coordinate was less than 275 pixels. After confirming that the crow movie did not provoke significantly different reaction in the subject, we conducted a cued conditioning experiment using seven crow vocalizations (Fig. 7c). Even to the naïve birds, the playback of crow calls triggered a marked behavioral response: upon hearing the sound, they typically looked around, turning their heads both left and right ^24^. This behavior was evidenced by an increase in head rotations, either clockwise or counterclockwise, of 10° or more (Fig. 7d). Concurrently, the birds remained relatively stationary, maintaining their position after the stimulus. During the conditioning sessions, we focused on determining whether the bird’s body-direction was directed towards the monitor (within ± 80°) or away form it (beyond ± 100°). Notably, we observed a reduced tendency for the subject to face the monitor in response to the crow movie presentation during the conditioning. This trend was evident in the subject’s reaction to the crow movie when its attention was captured within the initial 15 s (150 frame) while maintaining its position within a range of 10 pixels (approximately 12.56 mm) (Figs. 7f,g). The change in response became evident around the 6th sessions, conducted on the second day of conditioning, indicating the establishment of conditioned response to the stimulus. Following this conditioning phase, the bird’s reaction to the crow movie presentation changed, exhibiting an increased tendency to face away from the monitor (beyond ± 80°) during the initial 25 s of the movie presentation, even though the crow call was not presented (Fig. 7h). These changes were not observed in response to the Bengalese finch movies, which were not conditioned to crow calls. These observations suggest that our system is effective for longitudinal studies assessing learning-related changes in bird’s behavior.

**Fig. 7.**
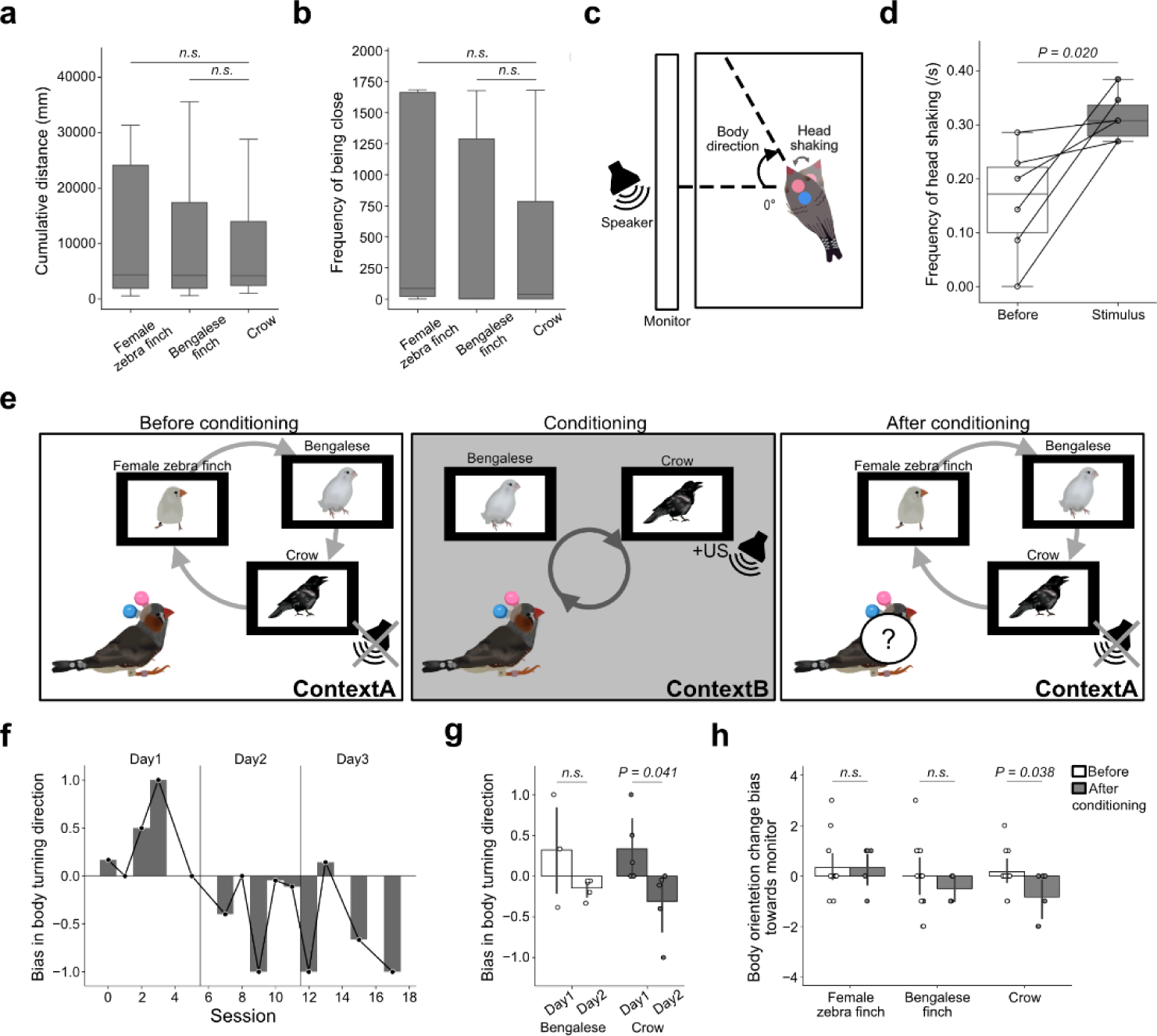
| Learning-dependent changes in behavioral reactions to stimuli. (**a**, **b**) Behavioral responses to the presentation of movies of zebra finch female, male Bengalese finch and crow to a bird naïve to crow. Cumulative distance (**a**) and frequency of being close to the monitor (**b**), during 3-min sessions. Tukey HSD test, 20 trials. (**c**) Diagram of the behavioral test with crow vocalization stimuli. (**d**) Frequency of head shaking before and during playbacks of crow vocalizations. Paired *t*-test, 6 trials. (**e**) Experimental diagram for analyzing behavioral reactions to movies before, during, and after cued conditioning. (**f**, **g**) Body turning direction towards versus away from the monitor, observed immediately following the crow movie (CS) observation in the conditioning session. The change in body turning direction bias in each session (**f**), and the summarized data from days 1 and 2 (**g**). (**h**) The change in body orientation before and after conditioning. The behavioral response to the movies was analyzed during the initial 25 s of the presentation. A change in body orientation was defined as the initial direction in which the bird consistently orientated for > 1 s (10 frames) following a directional change occurring within 5 s after observing the movie. These values were standardized based on their total and weighted by the frequency of observation that resulted in orientation change. Independent-sample *t*-test were used for comparisons in (**g**) and (**h**). Number of trials: Bengalese-day-1, 4; Bengalese-day-2, 5; crow-day-1, 5; crow-day-2, 5 in (**g**); female zebra finch-before; 15; female zebra finch-after, 6; Bengalese-before, 14; Bengalese-after, 4; crow-before; 12; crow-after, 6 in (**h**). n.s. not significant. Boxplots display the median and the interquartile range ± 95% CI. Bar graphs in (**g**) and (**h**) display the mean ± 95% CI.

## Discussion

The aim of this study was to develop an easy-to-use and cost-efficient method for analyzing complex animal behavior during social interactions. To achieve this, we adopted a marker-based motion capture system using just a single camera within a small arena. This contrasts to the typical motion capture system, which necessitates large arenas and rely on calibrated camera arrays ^7,13^. While our system currently captures behavior in two dimensions due to its single-camera setup, it successfully captured complex behaviors during social interactions. The inclusion of additional camera or the introduction of monocular camera distance estimation could enable three-dimensional posture tracking in the future. One of the drawbacks of using marker-based tracking is the difficulties in stably affixing the markers to the subject’s body. We addressed this issue by attaching screws directly to the bone. This approach ensures that the marker remain securely in the same location on the subject’s body for long time, facilitating longitudinal analysis of movement changes over time. Furthermore, consistent marker placement and detection accuracy across multiple individuals enables a precise measurement of individual differences in behavioral responses ^25^.

In this study, we developed a computer algorithm involving multiple processes to ensure the robust detection of markers and body direction. While HSV-color feature extraction-based color marker tracking itself comprises difficulty in tracking without incorporating noise or accidentally capturing off-target areas, we overcame this issue by implementing the MoG background subtraction method. Our motion capture system demonstrated its efficiency in a variety of environments and even with multiple subjects. This suggests it holds advantages over marker-less tracking systems such as DeepLabCut ^26^ in terms of versability and reliability. A common issue with such marker-less method is the difficulty in discriminating multiple subject with similar body features, which we addressed by utilizing Optical Flow. With this method, we successfully analyzed each individual’s behavior during social interactions. As a potential future development, our algorithm could be applied to track any distinctly colored head mounted objects, such as the Miniscope or the electrodes for neural activity recordings ^27,28^. One caveat of our system is the effect of markers. It have been reported that zebra finches have color preferences for artificially attached leg band or feather crest on their head ^29,30^. To minimize this effect, our analyses were conducted after sufficient acclimation of the markers on their head, with consistent usage of maker colors. However, further in-depth analysis of birds’ behavior during social interaction, comparing results obtained from marker-based and marker-less analysis, might provide improvements or new developments in behavioral analysis tools.

While motion analysis alone cannot fully reveal how animals perceive visual stimuli, it is noteworthy that birds have a limited range of eye movement, often confined to just a few degrees ^9,31^. Some birds including the zebra finch, have a single fovea ^18^. Consequently, their sight can largely be inferred from their head direction. Prior research has suggested that many birds predominantly employ monocular fixation when focusing on the attention catching objects ^17,18^. Through the sight analysis, we discerned a non-uniform preference in their visual field usage, particularly favoring the ± 60° direction from the head direction (Fig. 2g). We hypothesize that this orientation is optimal for zebra finch’s sight. Indeed, this direction aligns with the bird’s fovea, positioned at 65°, and coincides with the fixation angle in monocular vision, which was revealed to be around 50–65° from the beak angle ^18^. Hence, it can be concluded that our motion capture system accurately recorded the probable line of sight of the subjects, and provides an objective method to infer their attention.

Virtual signals offer a valuable means for conducting research in animal communication, as their contents can be easily manipulated, and identical signals can be utilized in repeated trials, thereby enhancing the reproducibility of research findings. Consequently, many studies have employed virtual signals, such as movies and robotic models, to explore the requirement of social information in the vocal communication of songbirds ^32–35^. For utilizing virtual reality signals in animal research, it is necessary to determine whether these virtual signals are perceived in manner similar to live signals. Our motion capture analysis can be utilized for such purpose. In this study, we observed differential behavior according the signal contents, even for virtual signals. However, we also detected differences in behavior between live signal and virtual signals. This may be derived from quality of signals, as live signals typically include more information than what is provided by virtual signals ^36,37^. The difference in the quality of information between live and virtual in our study might include, sounds, olfactory signals, textual information like wind vibration, and contingent responses ^38,39^. These features should be improved in the development of virtual signals that can truly substitute for live stimuli.

In conclusion, we have developed an efficient and cost-effective motion capture system for songbirds. By use of this system we can robustly analyze the body trajectory and the body and head direction on two-dimension arena, simultaneously for multiple subjects. These features can be used to infer their sight and attentions. This system will aid performing quantitative and precise analysis of songbird behavior during communication.

## Acknowledgments

We thank the members of the Abe Lab at Tohoku University for help and fruitful suggestions. This study was funded by Tohoku University Research Program “Frontier Research in Duo” No.2101, JSPS/MEXT KAKENHI 23K18252, 22H05482, 21K19424, 19H04893 to KA.

## Author Contributions

MF and KA conceived and initiated the project and wrote the manuscript. MF developed the methodology, performed all the experiment, data analysis, and prepared the initial manuscript. All authors discussed and commented on the manuscript.

## Data availability

Data are provided upon reasonable request for the corresponding author.

## Methods

### Animal treatment

The care and experimental manipulation of animals used in this study were reviewed and approved by the institutional animal care and use committee of Tohoku University. All experiments and maintenance were performed following relevant guidelines and regulations ^40^. Finches were purchased from Asada Chojyu and kept in our aviary under a 14-hour light / 10-hour dark cycle (daytime: 8:00−22:00); water and food were given *ad libitum*.

### Surgery

Adult zebra finches (*Taeniopygia guttata*) were anesthetized with medetomidine-midazolam-butorphanol mixture (medetomidine 30 mg/mL, All Japan pharma; midazolam 30 mg/mL, Astellas Pharma; butorphanol tartrate 500 mg/mL Meiji Seika Pharma; NaCl 118 mM Fujifilm-Wako; 100 mL per bird). Then, two small plastic screw (0.04 × 2 g) were firmly cemented to the cranial bone along with the rostral-caudal axis of midline. After recovery and sufficient affiliation to the screws, light-weight plastic color markers (two from red, blue, green, purple) were affixed to the screws via a small plastic nut glued on a hollow plastic ball (total 0.59 g) during behavioral analysis. Subjects were habituated enough to the head markers by the experiments more than two days after surgery. They were isolated and habituated to each new experimental environment from at least one day before the test. Behavioral recordings were started from 9:30 a.m. and markers were attached at 8:00 a.m. In an experiment for evaluation, a plastic color marker was glued on their back skin using Alon-alpha (Toagosei) after removing feathers. Subjects were put into a metal mesh cage (36.5 × 25.5 × 26 cm) situated within a soundproof chamber (inside dimension: 68 × 50 × 40 cm). Movies were recorded at 10 frames per seconds (fps) at 480 × 640 resolution by a single 180° fisheye camera (ELP, USBFHD06H-L180) mounted on the chamber’s ceiling.

### Motion capture system

We created a *Python* (v3.8.8) based architecture for marker tracking and estimation of bird location and body direction. The code implementing the MCFBM and documentation are available on GitHub (https://github.com/MizukiFujibayashi/MCFBM). To achieve robust marker tracking, we devised a computer algorithm that utilizes HSV (hue, saturation, value) color feature extraction with the Mixture-of-Gaussians (MoG) background subtraction (MoGBS) method for each frame. In some application, silhouette detection was performed using Optical Flow methods. The implement of MoG based background subtraction allows for robust detection of markers, as relying solely on the HSV color feature extraction tends to yield poor marker detection over the background noise. With movie recording at 10 fps, the MoGBS algorithm typically identified the body, even during flight. To detect the markers, all MoGBS contour lines were bolded (3 pixels), then blurred with 5 × 5 filter. Subsequently, contours of pixels with values greater than 0 were extracted. These contours were filtered based on their area, with a threshold of 600 pixels. The three contours closest to the center of the most recent marker positions (reference point) were selected. As a result, an image that possibly include the bird silhouette with minimal noises was obtained. After this preparation phase, HSV color feature extraction was applied for each color marker detection. From this phase, the application scope was restricted to a rectangular area centered on the reference point, with an expanding range corresponding to the number of marker-lost frames since the last tracked frame. Blurred frames were used for HSV color feature extraction to increase the robustness of marker detection, as we encountered cases where the marker was recorded on movies as appearing to be segmented by the net on the cage roof. Along with the contour selection by area, circularity, radius, saturation, and the distance from reference point, HSV based contours were selected by the including ratio of MoGBS based bird-like silhouette. If the maximum ratio of those HSV-based candidates was smaller than 10%, contours from the overlapping image between the HSV masked image and MoGBS were added to the candidates, and the same selection algorithm was applied. The reference point of the first frame was manually defined, and marker detection relied solely on HSV color feature extraction when no MoGBS-based bird-like contours were detected. Bird’s body-location and body-direction was tracked from the estimated bird silhouette. Different algorithms were applied to detect silhouette for single and multiple subjects, as detailed in below.

#### Silhouette detection for a single bird

The algorithm was applied within a rectangular scope centered on the marker positions calculated by the marker tracking system (reference point). Contours were extracted from a MoGBS image after blurring (5 × 5). MoGBS contours were pre-filtered if they occupied 40% of the circle within an average bird size (50 pixels) from the reference point, indicting that MoGBS was detecting most of the bird. From the contours with an area greater than 600 pixels, we selected the one that either encompassed the majority of the head marker or had its center closest to the reference point. This contour was chosen as the prime candidate to reflect the bird silhouette. If the prime candidate did not include any marker points and was far from the reference point, or if the circumradius exceeded that maximum bird size, then the watershed algorithm was applied. For the prime candidate that did not fulfill 60% of the fitted ellipse, particularly in case including the spiky noise, a morphological transformation was applied.

#### Silhouette detection for multiple subjects

As MoGBS alone cannot consistently track the bird’s silhouette, a combination of MoGBS and Optical Flow ^20^ was utilized in this algorithm. The algorithm was applied within a 147-pixel radius from the center of marker positions calculated by the HSV mediated marker tracking (reference point) for each bird. This algorithm includes both forward and backward applications. In the forward application, flow vectors for each pixel between previous and current frames were calculated using Optical Flow. Based on these flow vectors, the projection of bird silhouette in the previous frame was acquired, and morphological transformation was applied to fill the cracks. Additionally, the actual silhouette of the birds were obtained using contours extracted from the MoGBS image after blurring (5 × 5) and filtering by area with a threshold of 600 pixels. For each subject, we selected the contour that either included the majority of the head marker center or had a center closest to the reference point. If there was no common contour among the subjects, these were defined as the individual silhouette of each subjects. To obtain a less noisy silhouette, if the individual flow projection occupies more than half of each contour, we used the overwrapping area between the silhouette image and the flow projection. If there was any common contour, from the contours larger than 600 pixels in the flow projection image, we identified the one that included most of the head marker center points or whose center was the closest to the reference point for each subject. If no silhouette was detected by the MoGBS, algorithm relied solely on the flow projection image. In the backward algorithm, the same process with inverted flow vectors was applied to frames where no bird silhouette was detected in the forward phase.

#### Body center and direction estimation

The detected silhouette was fitted with an ellipse, and the center of this ellipse was used to determine the bird’s body-center. Of the four vectors extending from the center to the apices, one was designated as the indicator of the bird’s body direction. Body-direction estimation was skipped in fames where either the head-marker detection failed or the silhouette estimation was not reliable, to ensure a more consistent and accurate assessment. Once the body-direction vector was estimated, the subsequent frame utilized this information to select the body-direction vector from one of the two vectors on the major axis. The selection was based on which vector had a smaller angle relative to the previous vector. From the second frame, body-direction vector was defined as the vector from the center of ellipse at angle of rotated previous body-direction vector by the average degree changes within the latest 3 frames. In case where the posterior marker on the head fell outside of a circle centered on the ellipse center, with radius equal to a quarter of the semi-major axis, both the a semi-major and a semi-minor axes of the ellipse quarter that encompassed it was considered as the indicators of direction. Otherwise, two axes on the quarter that encompassed the widest angular range between the previous body-direction vector and the predicted direction vector were selected. From this candidate set of semi-major and semi-minor axis, the semi-major axis was typically chosen, except when the length ratio of the major to minor axis was less than 1.2. In such cases, it is plausible that the minor axis better represent the body direction. Such cases occurs occasionally, for instance, when the subject is spreading its wings. The candidate that exhibits a larger average of tilting bias of previous body direction vector and prediction body direction vector to itself was selected. For the first time of estimation after skipping for more than two frames or the very first estimation, different strategy was adopted with first frames in some range and the rest of frames. For the first 300 frames unless the posterior marker on head is outside of circle centered on the ellipse center with radius quarter of semi-major axis, two sets of body directions started with each vector stemming from the center to apexes of major axis were traced until body direction was estimated for 100 frame without losing successive two frames. If two sets of body directions were tracked for 100 frames, the set with smaller average value of average distance between selected apex and two head markers were choose. If the value was equal, same value of later half was compared. For the remaining cases, the semi-major axis on the ellipse quarter was selected as a candidate only when the posterior marker on head was outside of circle centered on the ellipse center with a radius equal to a quarter of the semi-major axis. This was excepted in case where the length ratio of major and minor axes was less than 1.2, or when the posterior marker was closer to the ellipse center, such as when the bird is turning its head backwards. In such cases, the semi-axis forming the smallest angle with the vector from the anterior to the posterior marker on the head was chosen. The subsequent body direction estimation followed the algorithm already described.

### Evaluation of motion capture system

For the evaluation of motion capture system, 52 files recorded from one male subject were used. These recordings are taken during live or virtual presentation of female and male zebra finch, and processed as described below.

#### Evaluation of the marker detection accuracy

To evaluate the marker detection for a single subject, we analyzed the proportion of the frames where the algorithm detected both anterior and posterior head markers within the expected distance, out of the total frames where the two markers were detected. Since the distance between the two head markers was expected to be stable, we calculated the proportion of instances where this distance was less than 15 pixels. The score ranged between 96.51% and 100%, with a mean value of 99.56%, and varied depending on the bird posture. For evaluating the marker detection for multiple subject tracking, we analyzed the proportion of the frames where the algorithm detected both anterior and posterior head markers within the expected distance, out of the total frames where the two markers were detected in both subjects. The score ranged between 95.38% and 100%, with a mean value of 98.64%. To further infer the tracking accuracy, we analyzed the conditional probability of the frames within the entire movie where the algorithm detected the anterior head marker, and where it detected both anterior and posterior head markers within a 15-pixel range. This accuracy value was estimated to 94.33 ± 2.90% during interaction and 98.93 ± 1.39% when the subject was alone.

#### Evaluation of Bird-center and body-direction accuracy

To assess the accuracy of bird-center and body-direction estimation, a plastic marker was attached to the subject’s back, and we compared its location and the direction of the vector from it to the posterior head marker. The measured distance between the marker on the back and the estimated center was 11.13 pixels ± 5.24 (mean ± s.d.). This distance between the actual marker on the back and the estimated center is expected to be a stable, with some fluctuation caused by bird’s posture. Therefore, we investigated the standard deviation, which ranged from 2.26 to 9.31 pixels, confirming the accuracy and efficiency of the body-location estimation. The average angle between the estimated body-direction vector and the vector from the actual marker on the back to the posterior head marker was 30.59° ± 33.21 (mean ± s.d.). To evaluate bird-center prediction for multiple subject, we analyzed the probability of the frames within the entire movie where the algorithm detected and calculated the position of the bird-center.

### Behavioral analysis

#### Stimulus procedure

Stimuli, both live and virtual format, were presented for 3 min with randomized interval ranging from 8–10 min, and the subject’s behavior was recorded for 3 min before and during the presentation in all experiments. Stimuli were presented in random order for each session. For live stimuli presentation, the subject bird was kept in cage where an empty cage was placed next to it. Then, cages either containing a zebra finch or empty were swiftly exchanged with the empty cage. A transparent acrylic board was placed on one side of these cages. The cages were put next to subject’s cage with acrylic sides facing the subject. For virtual stimuli presentation, movies without sounds were displayed on an LCD monitor (IPS-LCD, Kenowa JP-11.6-1080, resolution of 1,920 × 1,080, 11.6-inch screen). A back ground image (an empty cage image) was continuously presented on the monitor, and a frame made of cardboard was placed between the subject’s cage and the monitor to make the signals appear as if they were from actual neighbor.

#### Live or virtual female and male presentation

For the behavioral comparison between responses to live zebra finches, subjects were exposed to an adult female zebra finch, adult male zebra finch, or an empty cage during a single session. This procedure was repeated across five sessions. Subjects were acclimatized to each other by being kept in a single cages for several days before the tests. Then, all birds were isolated for one day before the test. The next day, for the behavioral comparison of female- and male-directed behavior in response to virtual signals, movies of the same female and male zebra finches and an empty cage were presented to the same subject in each session. This procedure was repeated across five sessions. A picture of empty cage was used as background image. *Marker-based detection of multiple subjects:* For experiments aimed at tracking multiple subjects within the same cage, two males and one female zebra finches were used. One male subject remained consistently in the experimental cage while the other male subject or the female subject were introduced in each session. A total of five sessions were performed for male-male and female-male interaction. Subjects were habituated to each other through cages for several days, and all birds were isolated one day before the experiment.

#### Live or virtual individual discrimination of male zebra finches

For comparing behavioral responses to live and virtual male zebra finches, four familiar adult male zebra finches and one unfamiliar adult male zebra finch were used. The familiar four male zebra finches had been kept in the same cage with the subject for approximately one month. The unfamiliar male finch had been isolated and had never seen the subject before the experiment. For the behavioral comparison of live and virtual male zebra finches, movies of those five male zebra finches and a movie of an empty cage were presented in one session. The picture of empty cage was used as back image throughout the procedure. Five sessions were repeated on a day. The next day, for the behavioral comparison of live male zebra finches, the corresponding four male zebra finches were presented in live format to the same subject in each session. Four sessions was repeated on the second day. This experiment involved one male bird (Male #4) whose whole body feather was white. As we found that the subject of behavioral analysis exhibited female-directed-like behaviors against that white male, we only analyzed the behavior towards other four males.

#### Color reduced female presentation

A female zebra finch was movie recorded against a black background. To modify color information of the bird in each frame, the bird’s foreground was semi-automatically clipped by Canny Edge Detection from the absolute value difference from background image. To create color-modified movies, color modified clipped-bird images were embedded onto the background image. In addition to original 8-bit movie (original), we prepared clipped-bird embedded movie without color reduction (clipped), movies with color reduction to 4, 2, and 1 bit, monochrome movie (gray), and pixel shuffled movie (shuffled). Original movie, modified movies, and movies showcasing an empty cage were presented more than seven times during two-days of experiment. On the first day, modified movies and movies of an empty cage were presented in a single session in random order, and these sessions were repeated three times. Subsequently, original movies, modified movies without color reduction, and enlarged versions of the modified movie without color reduction were presented in a single session in random order, and this sessions were repeated for three times. On the second day, original movies, modified movies, and movies of an empty cage were again presented in a single session in random order, and this sessions were repeated four times starting from around 10:43 a.m. The behavioral parameters used for PCA analysis were: ’cumulative moving distance’, ’number of sight line crossing to the presented area’, ’number of active sight’, ’ number of head movements when the angular change between successive frames briefly surpassed 15.8° (value corresponding to the top 20%)’, ’number of body movements when the velocity briefly surpassed 2.782 pixels (value corresponding to the top 20%)’, ’average duration of active sight’, ’average duration of body moving instances where the velocity surpassing 1.29 pixels (value corresponding to the top 50%)’, ’ average duration of head moving instances where the rotational speed surpassing 4.53° (value corresponding to the top 50%)’, ’average duration of being close to the presented area where the x-coordinate is less than 275 pixels’, ’average duration of body directing to the presented area’, ’average duration of head directing to the presented area’, and ’deviation in the ratio of frames where the presented area is projecting to subject’s forward while the subject remains still without moving its head and body’ and ’deviation in the ratio of frames where the presented area is projecting to subject’s forward while the subject moves its body center’.

#### Assessing the Behavioral change during learning

We used audio files of crow calls captured form movie of crows (*Corvus macrorhynchos*) recorded in a local zoo (Asahiyama Zoo, Asahikawa, Japan). To assess the behavioral response of bird naïve to crow vocalizations, a total of seven crow calls (1–6 s in length) recorded from two crows were presented once at the beginning of a 3-min presentation period. The session was not repeated, and each vocalization was played once. Due to a program failure, one vocalization file was not played, so behaviors during the six playbacks were analyzed. To assess behavioral changes in response to the change in salience of the stimuli, we conducted a conditioning procedure and monitored the behavioral changes before, during, and after the conditioning. Movies of female zebra finches, male Bengalese finches (*Lonchura striata* domestica), two crows, and an empty cage were used as stimuli. The background image, showing a cage with a perch but no bird, was taken from the crow movie file and was continuously presented on the monitor throughout the procedures. Before and after the conditioning, four movies were presented to the subject in one session, and this session was repeated ten times a day. After two days of movie presentations, conditioning to crow movie was carried out. The conditioning process was conducted in a new chamber that had not been used for other tests (context B). In this chamber, the cage and monitors were arranged similarly, but the wallpaper was changed to provide a spatial cue for a different context. During the conditioning, the identical crow movies were used as the conditioned stimulus (CS), and seven crow vocalizations were used as the unconditioned stimulus (US). Six out of seven vocalizations were confirmed to evoke behavioral change described above. For each conditioning session, two randomly selected US were played at the timing of 20 s and 2 min 20 s from the start of 3-min CS presentation. The USs were played from a speaker (JBL, JBLPEBBLESBLKJN) placed behind the monitor. The Bengalese finch movie was presented to assess the effect of habituation resulting from the repeated presentations of the same signal. The Bengalese finch-movie, or the crow-movie (CS), was exposed in random order for one session, and six sessions were conducted each day. The subject was conditioned for three successive days. The combinations and orders of the US were randomized from the perspective of both the day and the entire three-day period in order to prevent predictive responses to the stimulus. Each US stimulus was played no more than once in one day. The feeding and water trays were removed at 9:00 a.m. and returned about one hour after the end of each day’s conditioning session. The day after conditioning, movies of a female zebra finch, a male Bengalese finch, two crows, and the empty cage were presented for one day. The analysis of the bird’s direction before and after conditioning during CS presentation was conducted only when the subject was stationery, observing the monitor for more than three frames (∼0.3 s) without moving its head (<5°) or body (<10 pixels), and turned the body direction within the following 50 frames (∼5 s). This criterion was necessary to exclude instances where the subject might not have noticed the appearance of the CS. The ratio of body’s directional change towards the monitor versus the opposite side was standardized based on their total occurrences. Additionally, this standardized difference was weighted by the frequency of observations to account for the degree of attention. This analysis differs from that applied during conditioning, as we hypothesized that the ongoing procedural context could influence bird behavior.

### Statistics

The alpha score of 0.05 was used to reject the null hypothesis. Independent sample *t*-test and paired-*t*-test were used for the comparison of two data sets. Tukey HSD-test was employed for multiple comparisons. All statistical analyses were conducted using Python.

